# Pesticide exposure and adverse health effects associated with farm work in Northern Thailand

**DOI:** 10.1101/549618

**Authors:** Chanese A. Forté, Justin Colacino, Katelyn Polemi, Andrea Guytingco, Nicholas J. Peraino, Siripond Jindaphong, Tharinya Kaviya, Judy Westrick, Rick Neitzel, Kowit Nambunmee

## Abstract

**Objectives:** To assess pesticide exposure and understand the resultant health effects of agricultural workers in Northern Thailand.

**Design:** Cross-sectional study.

**Setting:** The entirety of this study was completed in Chiang Rai, Thailand, at Mae Fah Luang University, Chiang Rai Prachanukroh hospital, the village leader’s home, and the community center.

**Participants:** 97 men between the ages of 22-76 years of age; 70 were conventional farmwokers and 27 did not report any prior farm or pesticide spraying.

**Primary Outcome Measures:** We quantified exposure to pesticides including chlorpyrifos, methomyl, and metalaxyl, by air sampling and liquid chromatography/mass spectrometry. We estimated differences in self-reported health outcomes, complete blood counts, cholinesterase activity, and serum/urine calcium and creatinine concentrations at baseline between farmworkers and comparison workers, and after pesticide spraying in farmworkers only.

**Secondary Outcome Measures:** We quantified

**Results:** None of the farmworkers wore standardized PPE for the concentrated chemicals they were working with. Methomyl, ethyl chlorpyrifos, and metalaxyl were detected air samples in varying amounts. When it came to reporting confidence in the ability to handle personal problems, only 43% of farmworkers reported feeling confident; which reflects higher stress levels in comparison to 78% of comparison workers (p = 0.028). Farmworkers also had significantly lower monocyte counts (p=0.01), serum calcium (p=0.01), red blood count (p=0.01), white blood cell count (p=0.04), and butyrylcholinesterase activity (p<0.0001), relative to comparison workers. After adjusting for BMI, age, and smoking, methomyl air concentrations were associated with a decrease in farmworker acetylcholinesterase activity (beta= −0.327, p =0.016).

**Conclusions:** Farmworkers’ blood analytes, acetylcholinesterase, and self-reported symptoms differed from comparison workers. Improving PPE use presents a likely route for preventive intervention in this population.

Strengths and Limitations:

- The US Occupational Safety and Health Administration (OSHA) generally recommends testing for baseline cholinesterase levels after not working with organophosphates for at least 30 days(1). However, this was not capable for our study, and baseline cholinesterase measurements may not actually represent a true baseline measurement due to their overlapping work schedules
- This pilot study did not take multiple baseline measurements, and the one baseline that was taken was likely taken before the OSHA recommended guideline of 30 days since pesticide use.
- Our study also focused on workplace sampling at a time when the specific farm of interest was expected to be spraying chlorpyrifos, therefore the study results show an over-representation of chlorpyrifos.
- This is the first study of its type that took a mixed-methods approach using survey, biomarker, and workplace observation data to analyze farmworker pesticide health effects in comparison to other workers in Northern Thailand.
- This pilot study is one of the larger studies on farmworker chemical exposures in Thailand. These data can inform the methods for future global occupational health research on farmworkers.
- This study is very generalizable to farmworkers in LMIC and Thailand.

## Introduction

Agriculture is a major contributor to Thailand’s economy(2,3). Among surveyed farmers residing in Northern Thailand, most (97%) reported using pesticides on their crops (4–7). Over 93% of agriculture workers in Thailand work through the informal sector and are not defined as employees under the Labor Protection Act; thus, these workers are exempt from numerous safety laws surrounding labor and social security rights(8,9). Thailand ranked third out of 15 Asian countries for pesticide use per unit area and fourth in annual pesticide use(3). Although there have been increases in organic food consumption in Thai markets, there is no evident reduction in pesticide use(3,8). In fact, pesticide use in Thailand has increased from 110,000 tons in 2007 to roughly 172,000 tons in 2013(3). For the pesticides being imported into Thailand, a third are considered extremely, highly, or moderately hazardous based on the World Health Organization’s (WHO) hazard categories(3,10).

Pesticide pollution in the environment is associated with poisonings, oxidative stress, neurological dysfunction, birth defects, reproductive disorders, metabolism disorders such as diabetes mellitus, and cancers including colorectal cancer, prostate, and non-Hodgkin lymphoma(9,11–15). Although Thailand does not have a poisonings registry, a poison center located in Bangkok recorded more than 15,000 patients over a three-year period, 42% of whom had poisonings related to pesticide exposure(9). The pesticides most associated with these poisonings were insecticides: Carbamates, organophosphates, and pyrethroids(9). Research is lacking on low and middle income countries (LMIC) pesticide exposure and related health outcomes(16).

This study was motivated by a group of contract farmers residing in Wiang Pa Pao, Chiang Rai Province. These farmers reported concerns about their health related to spraying pesticides to Chiang Rai Prachanukroh hospital employees. Our project was launched in response to this concern. The purpose of this project was to assess the pesticide exposure of these farmworkers in Northern Thailand and to understand the resultant health effects when compared to workers residing in a non-agricultural area (Chiang Rai, Thailand). This project adopted a mixed-methods approach during assessment of pesticide exposure and the resulting health effects including: 1) personal air sampling, 2) biomarker sampling, and 3) perceived exposure and health effects assessed by questionnaire.

## Materials and Methods

The STROBE cohort reporting guidelines and checklist were completed in the creationv of this report.

### Study Population

All participants provided oral consent to participate in the study which was noted by the researcher administering the survey in the first step of study participation. The study protocol was approved by the Institutional Review Boards of Mae Fah Luang University and the University of Michigan (UM Submission ID: HUM00128091/CR00068767). All study participants were 21 years and older, male, and resided in Thailand. Participants were recruited in Chiang Rai, Thailand, using a recruitment script in Thai administered by health volunteers— laypersons trained through the Chiang Rai Prachanukroh hospital on patient care and interactions. Comparison workers were recruited through word of mouth at Mae Fah Luang University (MFU), with a focus on older males in work fields unrelated to agriculture with no commercial pesticide spraying experience. Farmworkers and comparison workers were recruited between July 1^st^, 2016 and September 15^th^, 2016. Overall, there were 123 study participants recruited and examined for eligibility. Twenty-seven comparison workers and 70 farmworkers were retained in the study who met the following inclusion criteria: were 21 years or older, male, residing in Northern Thailand and completed all follow-up.

### Sample Collection

All participants completed the consent form by oral confirmation due to literacy differences. Conventional farmworkers consented to allow work observation, active air sampling, and pre- and post-spray urine and blood samples to be taken. Each study participant also completed a researcher-administered survey. Participants received 600 Thai baht for participation and completion of the study.

We quantified exposure and potential adverse health effects among workers by selfreport, biological, and environmental sampling. A 69-item occupational health questionnaire was translated into Thai from English by MFU researchers. The questionnaire was based on a previous study of worker health and occupational noise(17). The survey was administered to study participants at the local community center and took roughly 35 to 45 minutes to complete. Questionnaire data were collected from July 2017 to the end of August 2017 (conventional farmworkers and comparison workers) and again in January 2018 (sample of comparison workers from Chiang Rai).

Nurses collected 10 mL of urine and 10 mL of blood from workers at the local hospital (Chiang Rai or Wiang Pa Pao). For farmworkers, blood and urine samples were collected within one week prior to pesticide spraying and again within 3 days after pesticide spraying. Whole blood samples were collected in red-top tubes with no anticoagulant, lavender-top tubes with EDTA, and green-top tubes with heparin.

### Blood and Urine Analysis

Urine creatinine, urinary calcium, serum creatinine, serum calcium, and complete blood counts were quantified by Meng Rai Laboratory in Chiang Rai. Acetylcholinesterase (AChE) and butyrylcholinesterase (BuChE) were analyzed using the Ellman method, from whole blood and serum, respectively(18). AChE was analyzed since it is directly related to AChE inhibition by the pesticides, and BuChE since it can be a sensitive biomarker of exposure to AChE inhibitors(19). Concentrations of the health biomarkers were compared between conventional farmworkers and comparison study participants, as well as within farmworkers before and after spraying.

### Air Sample Collection and Analysis

Personal pesticide air exposure levels were measured in conventional farmworkers only. GilAir Plus Personal Air Sampling Pumps by Sensidyne, Inc. were calibrated each day with the Gilibrator Calibrator before heading to the field. Air samples were collected during the time farmers were spraying pesticides (14 to 63 minutes) using XAD-2 sorbent tubes (SKC Ltd) based on the standard NIOSH method of 5600 at a flowrate of one liter per minute(20). Field blanks were collected by opening tubes away from the farms for a similar period of time; no air was drawn through the blanks, but they were otherwise handled identically to actual air samples.

The analysis of pesticide residues was completed by the Lumigen Instrument Center at Wayne State University and researchers were blinded to exposure or worker category. Laboratory controls from a spiked and blank filter were extracted each batch. A calibration check at 10 ng/mL and a solvent blank were run every 10 samples. If a check did not pass within 20% error, the entire 10 sample section was re-run. Calibration curves were prepared the same day per batch by comparing the concentration of and area ratio of analytes to deuterated surrogates. A linear regression was taken, and percent error calculated at each calibration point. If a point did not fall within 20% error of the predicted value, it was dropped from the end of the curve.

Air sampling tubes were extracted as either top portions or bottom portions. Top portions of the tubes contained the retainer ring, filter paper, foam pad, and XAD fill. Bottom portions contained the middle foam pad, and the bottom portion of XAD fill; the bottom foam was discarded. Portions were transferred to a 2 dram vial followed by 1960 μL of methanol and 40 μL of a solution containing 1 μg/mL each of deuterated surrogates. The vials were tightly capped and sonicated for 30 minutes, allowing the bath to heat naturally. Heating was found necessary to achieve equilibration of the surrogates and standards to the XAD fill and improve recoveries. Samples were centrifuged to settle any particulates and 900 μL of sample was diluted with 100 μL of water containing 100 mM ammonium formate, resulting in a final sample containing 10% water and 10 mM ammonium formate. Liquid chromatography mass spectroscopy (LCMS) analysis was completed using a Nexera-X2 ultra performance liquid chromatography (UPLC) with Shimadzu 8040 triple quadrupole mass analyzer. Chromatography was achieved using a Waters acquity UPLC Ethylene-Bridged Hybrid (BEH) C18 (1.2 μm, 2.1 x 50 mm) column eluting with 10 mM ammonium formate in water (mobile phase A) and 10 mM ammonium formate in methanol (mobile phase B).

### Worker Observations

MFU and UM student researchers observed the farmworkers during spraying activities and recorded worker practices for mixing, spraying, and storage of the pesticides on a worker observation form. This included information on the pump used, pump calibration, personal protective equipment (PPE) used, and notes on common practices. Detailed notes on PPE used such as item and material were noted by observers (e.g. gloves made of rubber, latex, or cloth). Work observations were not completed for comparison workers since they do not perform commercial pesticide spraying.

### Statistical Analyses

For questionnaire data, we calculated summary statistics of demographic data across the entire population. We first tested for crude differences across the measures between the conventional farmers and the comparison workers by t-test for continuous variables or by Fisher’s Exact test for categorical variables. Analysis of variance (ANOVA) and multivariate regression analyses were used to compare serum and urinary biomarker concentrations between farmworkers and comparison workers, adjusting for the potential confounders age, smoking status, alcohol consumption, and BMI. We also conducted multivariate linear regression analyses of the association between pesticide levels quantified in the personal air samples and serum and urinary biomarkers in the farmworkers collected after spraying, adjusting for the same covariates as described above. Any variables that were not normally distributed were log transformed prior to regression. SAS 9.4 was used to complete regressions and demographic tables, and R 3.5.1 was used to create figures and graphics.

### Data Statement

The de-identified data can be made available via Dropbox upon reasonable request and agreement to a memorandum of understanding of ethics.

## Results

Initially we recruited 73 farmworkers and 29 healthcare workers, and upon further recruitment we were able to secure more general comparison workers through word of mouth recruitment at Mae Fah Luang University. This new general comparison worker group was comprised of 27 men with similar mean age and education backgrounds as the farmworkers. Ultimately, the healthcare workers were overly female and had college educations and were dropped. The 27 general workers and the 70 male farmworkers were retained for this study.

Overall, farmworkers and comparison workers had similar demographics, except for BMI, which was significantly lower for farmworkers (Table 1). Farmworkers had a median age of 50 years with a range of 22 to 76 years of age, and comparison workers had a median age of 49 with a range of 39 to 68 years. Most of the workers were married, attended primary and secondary school, drank alcohol 1-3 days per week, and were current smokers (Table 1).

**Table 1.**
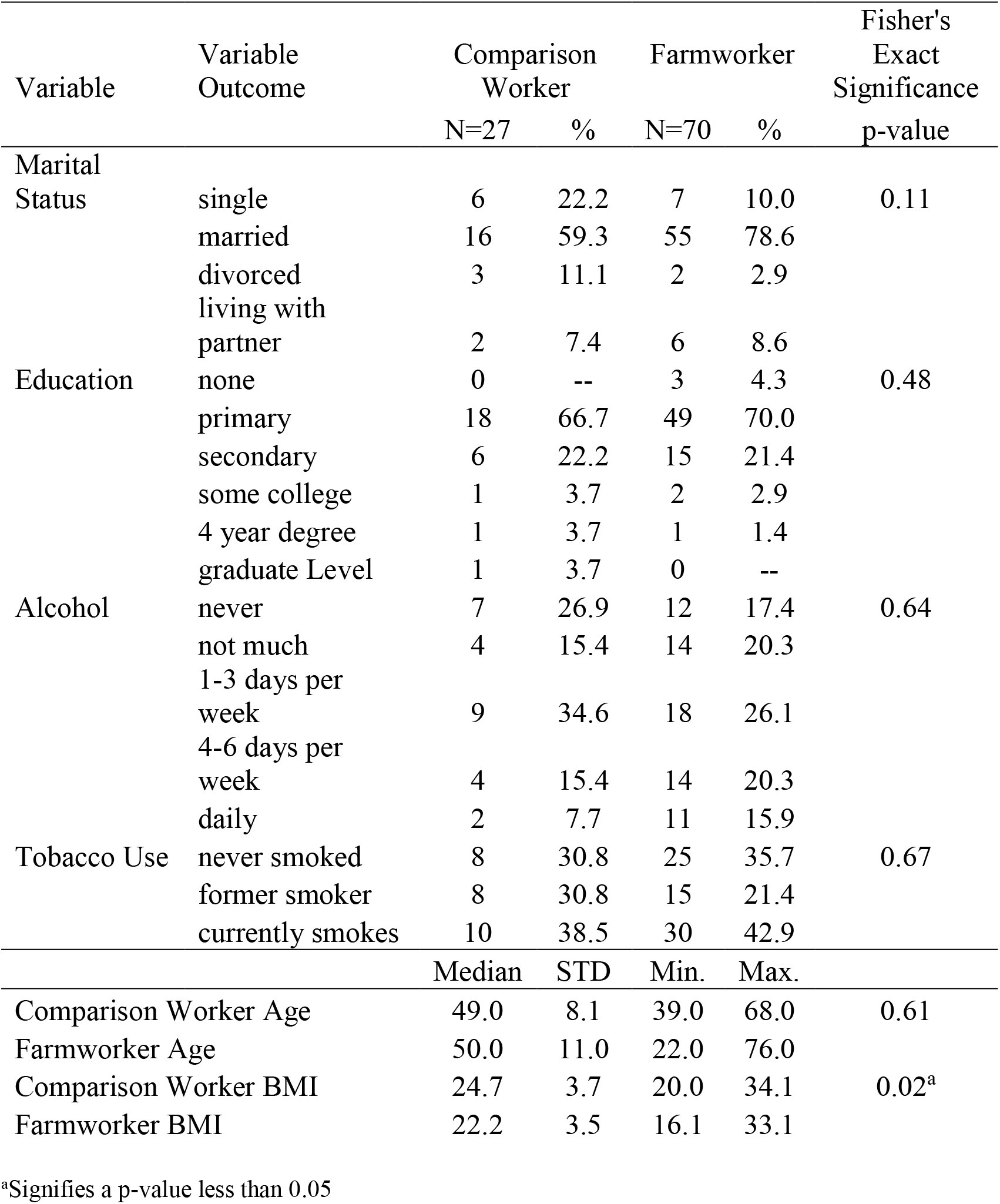
Study participant demographics, males only

Based on worker observations, none of the farmworkers wore chemical proof aprons, chemical proof gloves, or a respirator. The farmworkers did wear an item that we named ‘face hats’ which included both home-made and commercial versions. The home-made version was usually two towels or scarves (e.g. durags) stitched together to cover the head and face, and the commercial version was a canvas brimmed hat with a canvas face covering that could be removed. The use of gloves, long sleeve clothing, and any sort of clothes covering (e.g. rain poncho or plastic sheet) were usually used items with some damage and use was not consistent across workers.

With respect to self-reported health concerns, farmworkers and comparison workers selfreported symptom responses only statistically significantly differed for ‘dizziness’ and ‘shaking or trembling of hands’ (Supplemental Table 2). Supplemental Table 3 presents worker selfreported stress levels by worker category based on the Cohen’s Perceived Stress Scale. Overall, comparison workers more frequently reported stress in comparison to farmworkers, although not statistically significantly so. However, when it came to reporting confidence in ability to handle personal problems, only 43% of farmworkers reported feeling confident often in comparison to 78% of comparison workers (p-value = 0.03).

A comparison of cholinesterase activity between the two groups indicated that although there were some farmworkers with higher AChE activity, the AChE levels in farmworkers and comparison workers were not significantly different (p=0.20; Figure 1A). Comparison workers had higher BuChE activity levels compared to farmworkers (p<0.0001; Figure 1B). AChE and BuChE concentrations were not correlated (r:-0.09, p=0.35, 95% CI: −0.29, 0.10). Within farmworkers only, pre and post-spray activity of AChE (0.32, p=0.01) and BuChE (0.31, p =0.01) were correlated.

**Figure 1.**
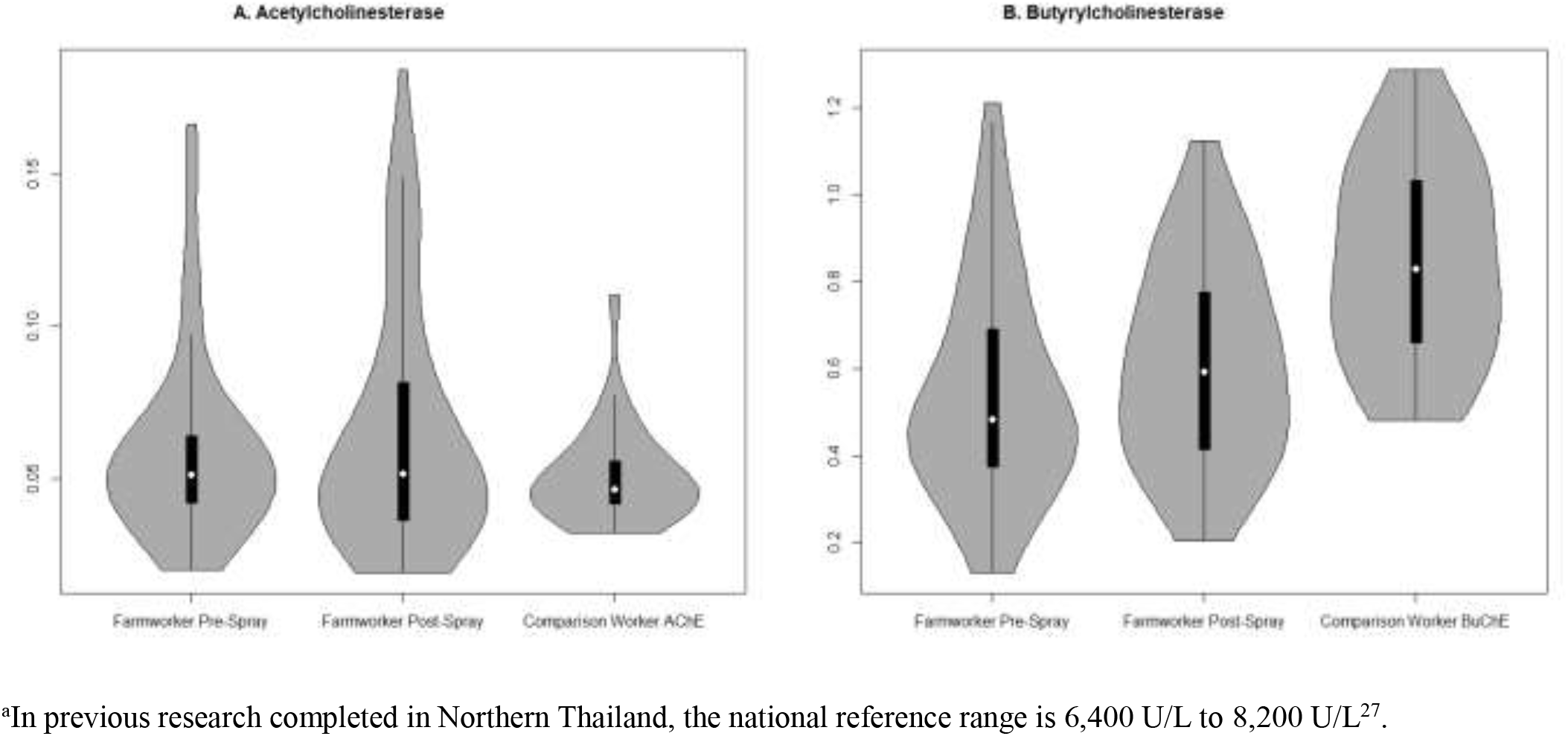
Distribution of acetyl- and butyrylcholinesterase activities in comparison workers (one time point only) and farmworkers (pre- and post-spray)^a^

Measurements of pesticide residues on air samples are displayed in Figure 2. Overall, ethyl chlorpyrifos and metalaxyl were detected at the highest frequency, while methomyl was less frequently detected. Most chlorantraniliprole (N=60) and methyl-chlorpyrifos (N=61) measurements were below the limit of detection. Ethyl chlorpyrifos, followed by metalaxyl and methomyl, was found in the highest concentrations in the air filter samples.

**Figure 2.**
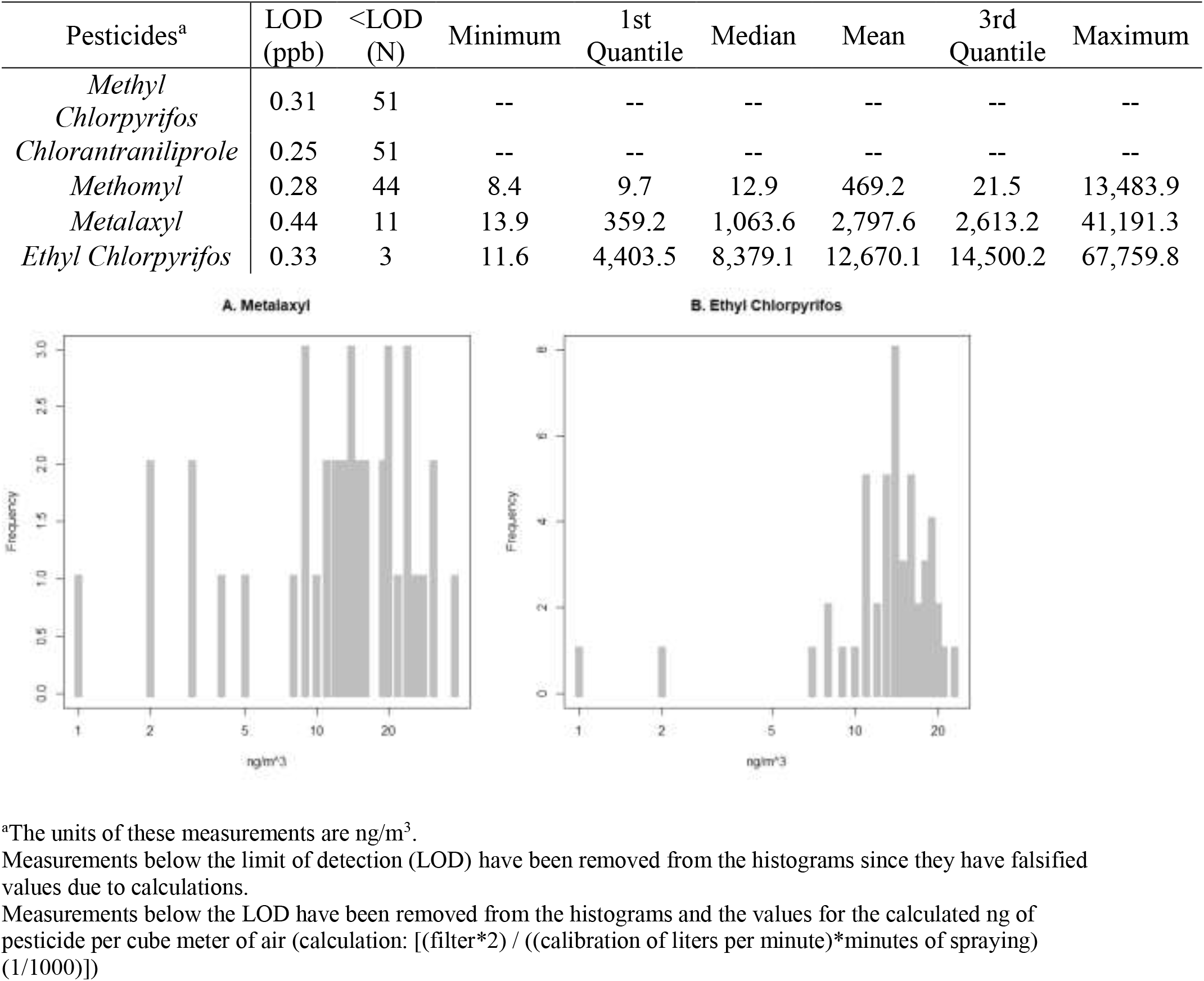
Pesticide measurements captured by personal air monitors on Northern Thailand farmworkers (N=51) with histograms showing distributions of detected values of (A) metalaxyl and (B) ethyl chlorpyrifos

Table 2 presents linear regression parameter estimates and confidence intervals comparing biomarker concentrations between farmworkers and comparison workers. Eosinophil (p=0.02), urine creatinine (p=0.03), and MCV (p=0.04) concentrations were statistically significantly higher in farmworkers than the comparison group. Monocytes (p=0.01), red blood cell counts (p=0.01), white blood cell count (p=0.04), and serum calcium (p=0.02) were statistically significantly lower in farmworkers than comparison workers.

**Table 2.** Linear regression analyses comparing biomarker concentrations in farmworkers and comparison workers

| Analytes | Units | Farmworkers=1 vs. Comparison=0 |  |  |  |  |  |  |  |
| --- | --- | --- | --- | --- | --- | --- | --- | --- | --- |
|  |  | Unadjusted Values |  |  |  | Adjusted Values |  |  |  |
| | | $\beta$ Estimate | p-value | 95% CI | | $\beta$ Estimate<sup>b</sup> | p-value | 95% CI | |
| log(Basophil) | % | 0.08 | 0.25 | 0.23 | -0.06 | 0.12 | 0.13 | -0.04 | 0.29 |
| log(Eosinophil) | % | 0.19 | 0.05 | 0.00 | 0.37 | 0.24<sup>a</sup> | 0.02 | 0.04 | 0.44 |
| log(Hemoglobin) | g/dl | -0.02 | 0.06 | -0.03 | 0.00 | -0.01 | 0.16 | -0.03 | 0.01 |
| log(Hematocrit) | % | -0.01 | 0.15 | -0.03 | 0.00 | -0.01 | 0.28 | -0.03 | 0.01 |
| log(Lymphocyte) | % | -0.04 | 0.25 | -0.10 | 0.03 | -0.03 | 0.41 | -0.10 | 0.04 |
| log(Mean corpuscular hemoglobin concentration (MCHC)) | pg | 0.00 | 0.37 | -0.01 | 0.00 | <0.01 | 0.53 | -0.01 | 0.01 |
| log(Mean Corpuscular Volume) | fl | 0.03 | 0.02 | 0.00 | 0.06 | 0.03<sup>a</sup> | 0.04 | <0.01 | 0.06 |
| Monocyte | % | -1.09 | 0.02 | -1.99 | -0.20 | -1.33<sup>a</sup> | 0.01 | -2.30 | -0.37 |
| Neutrophil | % | -0.48 | 0.87 | -6.49 | 5.53 | -1.34 | 0.69 | -8.08 | 5.39 |
| log(Plate Count) | cells/uL | -0.02 | 0.52 | -0.09 | 0.04 | -0.03 | 0.39 | -0.10 | 0.04 |
| log(Red Blood Cell (RBC) Count) | cells/uL | -0.04 | 0.00 | -0.08 | -0.01 | -0.04<sup>a</sup> | 0.01 | -0.07 | -0.01 |
| log(RBC Distribution Width (RDW)) | % | 0.01 | 0.15 | -0.01 | 0.03 | 0.01 | 0.19 | -0.01 | 0.03 |
| log(White Blood Cell Count) | cells/uL | -0.07 | 0.01 | -0.12 | -0.02 | -0.06<sup>a</sup> | 0.04 | -0.12 | <0.01 |
| Serum Calcium | mg/dl | -0.83 | 0.00 | -1.31 | -0.35 | -0.79<sup>a</sup> | <0.01 | -1.31 | -0.27 |
| log(Serum Creatinine) | mg/dl | -0.04 | 0.04 | -0.08 | 0.00 | -0.03 | 0.22 | -0.07 | 0.02 |
| log(Urine Calcium) | mg/dl | 0.13 | 0.12 | -0.03 | 0.30 | 0.14 | 0.14 | -0.05 | 0.32 |
| Urine Creatinine | mg/dl | 33.03 | 0.03 | 3.82 | 62.24 | 36.29<sup>a</sup> | 0.03 | 4.17 | 68.41 |
| log(Acetylcholinesterase) | U/L | 0.08 | 0.37 | -0.10 | 0.27 | 0.05 | 0.64 | -0.16 | 0.25 |
| log(Butyrylcholinesterase) | U/L | -0.48 | <0.01 | -0.68 | -0.29 | -0.50<sup>a</sup> | <0.01 | -0.70 | -0.29 |

<sup>a</sup>Signifies a p-value less than 0.05<sup>b</sup>These values were adjusted for age, former smoker status, current smoking status, alcohol use, and BMI.

Table 3 presents the linear regression beta estimates between biomarker concentrations and air sample pesticide content for farmworkers only. The ratio of pre- and post-spray AChE was significantly lower for increased concentrations of methomyl in air samples (p=0.02). Urinary creatinine and serum calcium were inversely associated with air sample concentrations of metalaxyl.

**Table 3.**
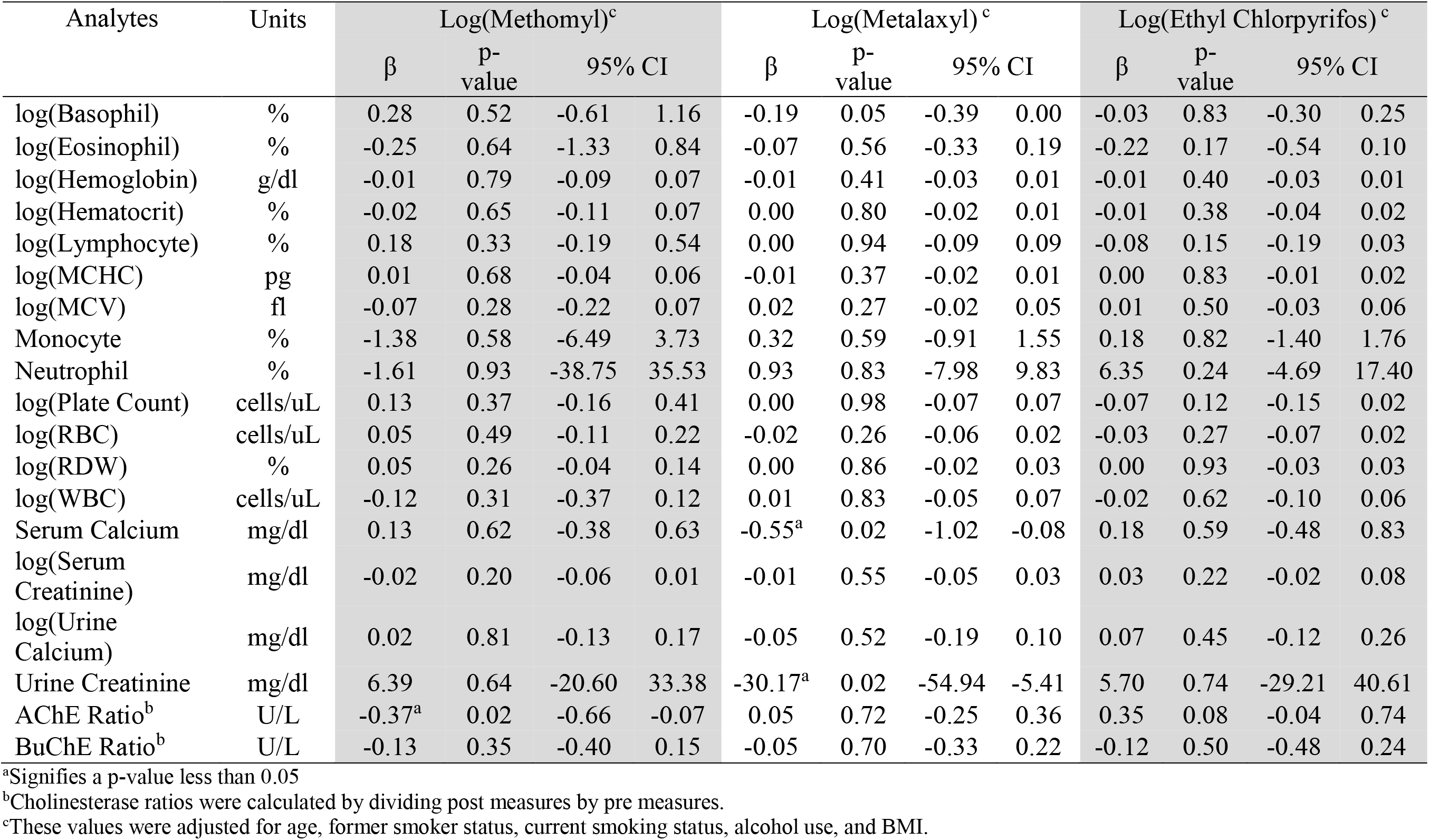
Linear regression models of association between blood analytes and pesticide air sample concentrations

## Discussion

This pilot study was designed to better characterize pesticide exposure and resultant health effects among farmworkers residing in Northern Thailand. We used a mixed-methods approach including collection of self-reported data, biological samples, and environmental air sampling. Farmworker and comparison worker demographics were not significantly different, with the exception of BMI, where farmworkers had significantly lower values most likely due to full work days doing physical labor.

Self-reported stress levels, such as the ability to handle personal problems, was reported less in farmworkers differing from comparison workers. Farmworkers and comparison workers responded significantly different on dizziness and in shaking or trembling of the hands. Biomarker concentrations also differed significantly between farmworkers and comparison workers. Farmworker concentations of urinary creatinine, MCV, Eosinophil were higher, whereas monocytes, WBC, serum calcium, and BuChE ratio were lower. Based on the regression analyses of only exposed farmworkers, serum calcium and urine creatinine were statistically significant among farmworkers exposed to metalaxyl. AChE ratio was also statistically lower for methomyl in the regression model controlling for age, BMI, current smoking status and former smoking status among farmworkers.

The most commonly used insecticides in Thailand are AChE inhibitors such as organophosphates and carbamates like chlorpyrifos and methomyl (3). In this study, 59 farmworkers self-reported using chlorpyrifos during pesticide spraying, 17 reported using metalaxyl, and 1 reported using methomyl. The analytic method utilized in this study quantified ethyl-chlorpyrifos, methyl-chlorpyrifos, metalaxyl, methomyl, and chlorantraniliprole. However, only ethyl chlorpyrifos, methomyl, and metalaxyl were detected on workplace air samples. Methomyl is a carbamate and insecticide and metalaxyl is a benzene compound used as a fungicide(21,22).

We observed workers during day 8 of edamame growing because they reported using chlorpyrifos during that time. This sampling timepoint likely resulted in our air samples not containing chemicals that are used at other times in the edamame growth cycle, based on farmworker self-reported chemical use and the spray schedule. Exposure to carbamates or OPPs can alter AChE levels from baseline(23–25). Considering the cross-sectional nature of this study and the frequency of pesticide spraying among our cohort, we cannot determine whether the cholinesterase biomarker results were due directly to the insecticides monitored or from another pesticide sprayed during another agriculture job or spraying another plant on the same farm.

On average, the farmworkers in our cohort worked 50.95-hour work weeks. The Labor Protection Act limit jobs that are deemed hazardous to human health a maximum of 42-hour work weeks. However, since 93% of all farmworkers in Thailand are informal sector workers, they are not working under the Labor Protection Act of 1998(8,9). Our cohort pilot study clearly demonstrates there are informal sector farmworkers working above the legal limit for hazardous jobs. For migrant farmworkers, this means they are likely working unhealthy hours and also lack access to the Thai healthcare system(26).

In a study by Sapbamrer and Nata (2017), rice farmers and non-farmer controls residing in Northern Thailand were interviewed and had blood samples taken to ascertain their overall pesticide exposure(27). When looking at self-reported health outcomes in our study, farmworkers did not have differences in respiratory symptoms relative to comparison workers. Farmworkers reported trembling in their hands less often in comparison to other workers, however with exposure to AChE inhibitors, we would expect farmworkers would report trembling more often.

Other studies have assessed the relationship between pesticide exposure and cholinesterase activity outside of Thailand. Strelitz et al. measured baseline to post-spraying whole blood and serum cholinesterase levels of 215 farmworkers from the Washington State Cholinesterase Monitoring Program. Ellman colorimetric enzymatic assays were used by two different laboratories to measure BuChE and AChE using the same methodology (Washington State Public Health Laboratories in 2006 and Pathology Associates Medical Laboratories in 2007 to 2011)(19). In this same study, cholinesterase depression was defined as 20% or greater decrease from baseline to post-exposure cholinesterase exposure(19). The authors found AChE and BuChE activity to be negatively correlated (−0.14, 95% CI: −0.27, −0.01)(19). Our study also found the correlation between AChE and BuChE to be weak, but the correlation was non-significantly positive (0.05, 95% CI: −0.199, 0.294). The ratio of pre-and post-spray AChE activity was also significant lower with increasing concentrations of methomyl in air samples. Others have reported BuChE activity as a more sensitive biomarker of exposure to cholinesterase inhibitors than AChE(19,28,29). In our study, BuChE activity was significantly lower in conventional farmworkers relative to comparison workers.

Garcia-Garcia et al. 2015 completed a similar study comparing 169 green house workers who were exposed to pesticides and controls who were unexposed to pesticides in southeastern Spain(30). Among this cohort, researchers controlled for sex, age, and BMI, but not smoking status. They found BuChE inhibition, RBC, MCV, and neutrophil levels significantly increased, whereas eosinophils significantly decreased and monocytes were not different(30). Our study assessed hematological parameters by measuring differences in complete blood counts between farmworkers and comparison workers, while controlling for BMI, age, former smoker status, current smoker status, worker category, and pesticide exposure levels. Serum calcium was statistically significant by both worker category and was also significantly different in farmworker. MCV, monocytes, RBC, eosinophils, urine creatinine and WBC were significantly different between our Northern Thailand farmworkers and comparison worker. In our study, farmworkers had a reduction, in RBC and neutrophils, whereas MCV increased (opposite of Garcia-Garcia et al.). Monocytes and eosinophils reduced and increased, respectively in our study and the Garcia-Garcia et al. study. In our study, an association existed between pesticide exposure and all of the aforementioned blood counts when looking at farmworkers and comparison workers.

Finally, a cohort study on farmworkers in China before and after pesticide spraying points to the effects of pesticides on complete blood count (CBC) having an effect that can be categorized as long term (3 years) or short term (10 days or less). Long term results include decreased white blood cell count. Short term results include as alterations in complete blood count, hepatic and renal functions, nerve conduction velocities, and on monocytes, hemoglobin and platelet counts(31). When comparing air sample measurements of methomyl, chlorpyrifos and metalaxyl, we did not identify differences in blood count analytes by chemical exposure, although we identified a range of differences when comparing farmworkers and comparison workers. Prior research has reported significant changes to hematological biochemistry as a result of pesticide exposures causing oxidative stress(30). Oxidative stress due to pesticide spraying has also been related to changes in CBC such as monocyotsis, leukocytosis, neutrophilia, and lymphocytopenia which were thought to also be closely related to patients in oxidative stress(30,32).

### Strengths and Limitations

Our study has some limitations. We focused on farmworkers who contracted to numerous farms, and therefore workers were likely exposed to other chemicals that were not quantified in this study. These farmworkers may have been exposed to chemicals that were not captured by this study because they sprayed pesticides on other farms and also sprayed other plants on the same farms. Due to this discrepancy, our farmworkers’ baseline cholinesterase measurements may not actually represent a true baseline measurement due to their overlapping work schedules. The US Occupational Safety and Health Administration (OSHA) generally recommends testing for baseline cholinesterase levels after not working with organophosphates for at least 30 days(1). OSHA also recommends taking a baseline measurement before and after organophosphate use (with at least a 30 day washout period for both)(1). This pilot study did not take multiple baseline measurements, and the one baseline that was taken was likely taken before the OSHA recommended guideline of 30 days since pesticide use. However, due to the crop production schedule and growing seasons in Northern Thailand, this population does not appear to ever experience thirty days of no pesticide use. Our study also focused on workplace sampling at a time when the specific farm of interest was expected to be spraying chlorpyrifos, therefore the study results show an over-representation of chlorpyrifos.

Overall, this is the first study of its type that took a mixed-methods approach using survey, biomarker, and workplace observation data to analyze farmworker pesticide health effects in comparison to other workers in Northern Thailand. Additionally, this pilot study is one of the larger studies on farmworker chemical exposures in Thailand. These data can inform the methods for future global occupational health research on farmworkers. Work observations also included a more detailed outline of PPE used by the farmworkers and could inform future studies on PPE in relation to pesticide exposure and preventive interventions. Overall, this study is very generalizable to farmworkers in LMIC and Thailand. This study will contribute to the building literature on farmworker pesticide exposure and resultant health outcomes.

## Acknowledgements

Participant recruitment, data collection and translation between English and Thai: Kanlaya Rattannasuwan, Anong Mungmueang, Aurai Channgoen, Rungthong Bunta, Rungtiwa N Lampang, Phornphan Saenchangmai, Kith Pakdeepong, Tharinya Kavuya, Nattaya Singkaew, Pimprapai Junsang, Phadungsuk Muanguk, Sratchaya Poolperm, Or-raya Onsongchan, and Wanwisa Kansa. We thank Dr. Kirsten Herold of the SPH Writing Lab, University of Michigan School of Public Health for her final review and editing.

## Patient and Public Involvement

This project was started based on farmworkers requesting a study be done. The local community, village leaders, and healthcare volunteers (laypersons with healthcare training) were imperative to the creation and completion of this research project. These community members were consulted and paid in kind for their expertise in identifying and communicating with stakeholders for the study, coordinating transportation, participant recruitment, and engagement, data collection, and translation between English and Thai (2 dialects of Thai included in this study). Additionally, the farmworkers also provided the research team with a generous gift of produce at the end of the project, and we thank them for their time, willingness to participate, and for the nourishment.

## Author Contribution

Kowit Nambunmee, Rick Neitzel, Justin Colacino, and Chanese Forte developed the study design. Katelyn Polemi completed the literature review alongside Chanese Forte. Chanese Forte, Andrea Guytinco, Siripond Jindhapong, and Tharinya Kaviya collected samples and administering the survey to farmworkers. Judy Westrick and Nick Peraino completed the air sample analyses and Nick Peraino drafted the methods section for the air samples. Chanese Forte analyzed the data under the supervision of Justin Colacino, Kowit Nambunmee and Rick Neitzel. Chanese Forte drafted the rest of the paper with all authors contributing with edits.

## Competing Interests

Authors have none to disclose.

No additional data.

## What this paper adds

1. What is already known about this subject?

- Agriculture is classified under the informal sector in Thailand which means many of the workers are not protected under worker safety laws.
- Thailand is the 3^rd^ highest ranked pesticide user out of 15 Asian countries and is increasingly importing more pesticides each year.
- Occupational pesticide exposure can cause of range of adverse health outcomes
2. What are the new findings?

- Methomyl, ethyl chlorpyrifos, and metalaxyl were detected in varying amounts in air monitor sampling.
- Conventional farmworkers also had significantly lower monocyte counts (p=0.01), serum calcium (p=0.01), red blood count (p=0.01), white blood cell count (p=0.04), and butyrylcholinesterase activity (p<0.0001), relative to comparison workers.
- After adjusting for BMI, age, and smoking, methomyl air monitor concentrations were associated with a decrease in farmworker acetylcholinesterase activity (beta= −0.327, p =0.016).
3. How might it impact on clinical practice in the foreseeable future?

- This study provides a new mixed methods approach to assessing worker exposure to chemicals, and therefore can highlight more adverse effects such as mental health, biomarkers, and PPE which were previously not possible in one study design.
- Farmworkers on average worked 50.96 hours per week which is above the Royal Thai Government Labor Protection Act of 1998 that limits work weeks to 42 hours.
- Farmworkers are exposed to high levels of pesticides without access to the appropriate PPE which could cause new company policies for PPE access or laws by the Royal Thai Government.

## Appendix A: Figures and Tables

**Supplemental Table 1.**
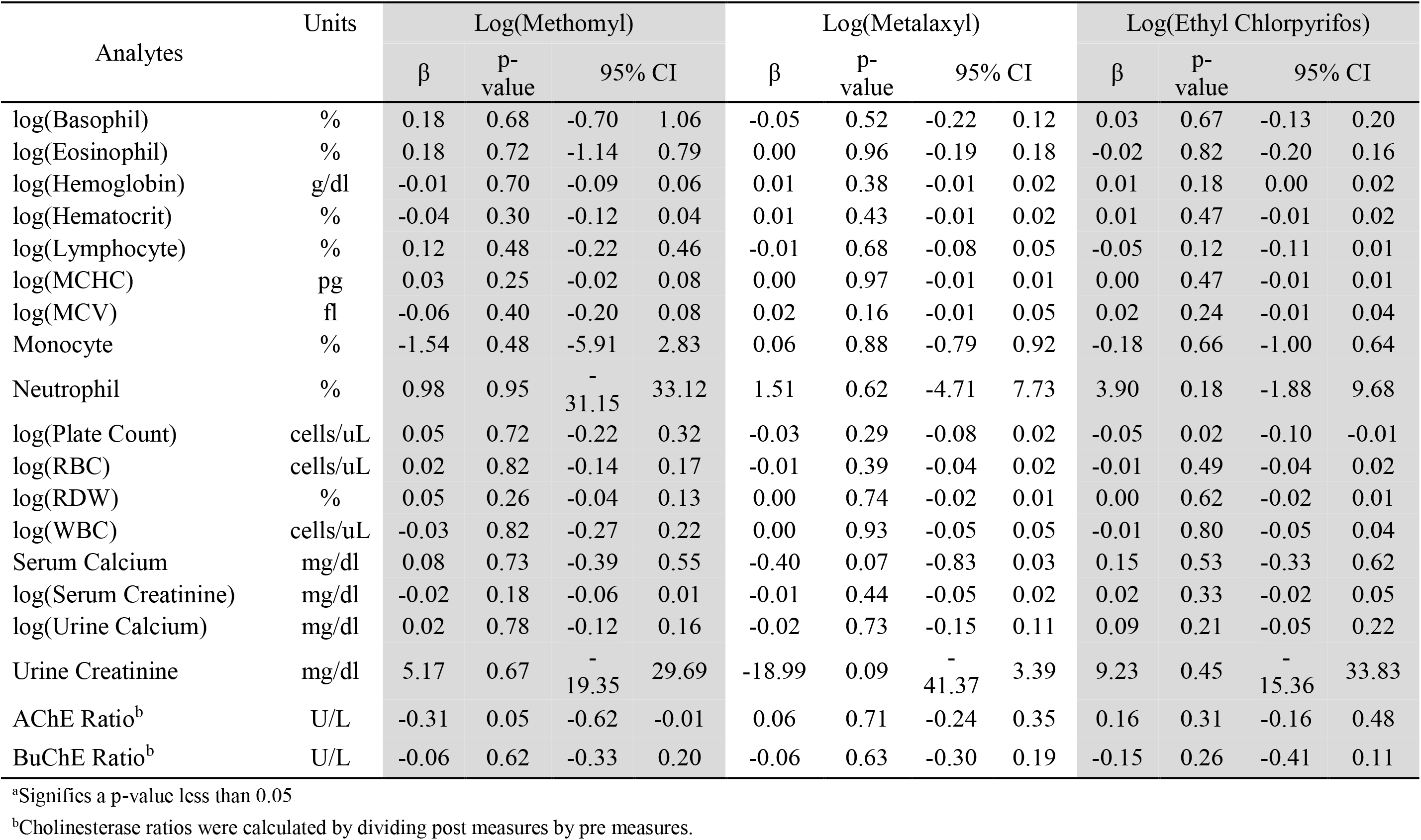
Unadjusted Linear regression models of association between blood analytes and pesticide air sample concentrations

**Supplemental Table 2.**
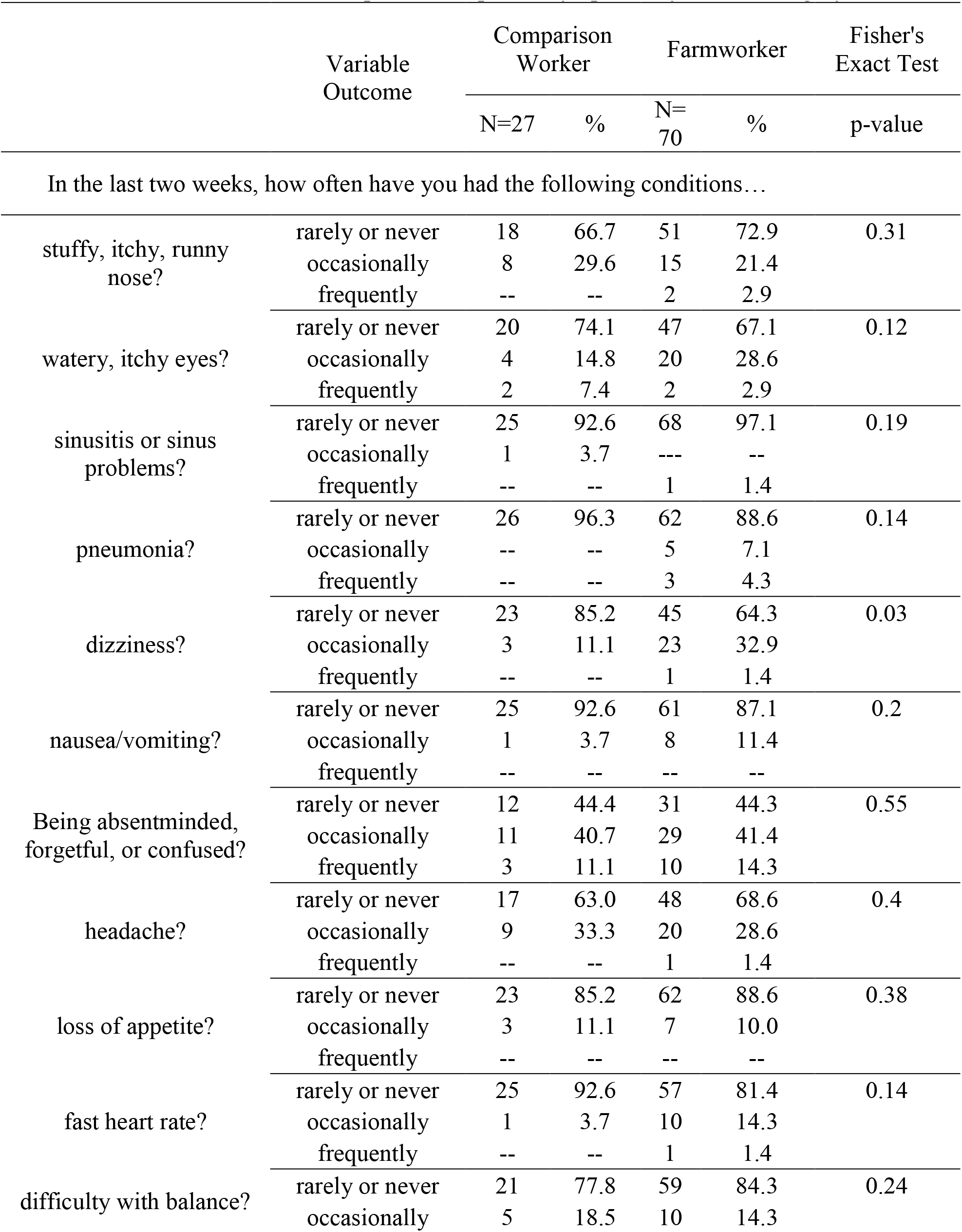
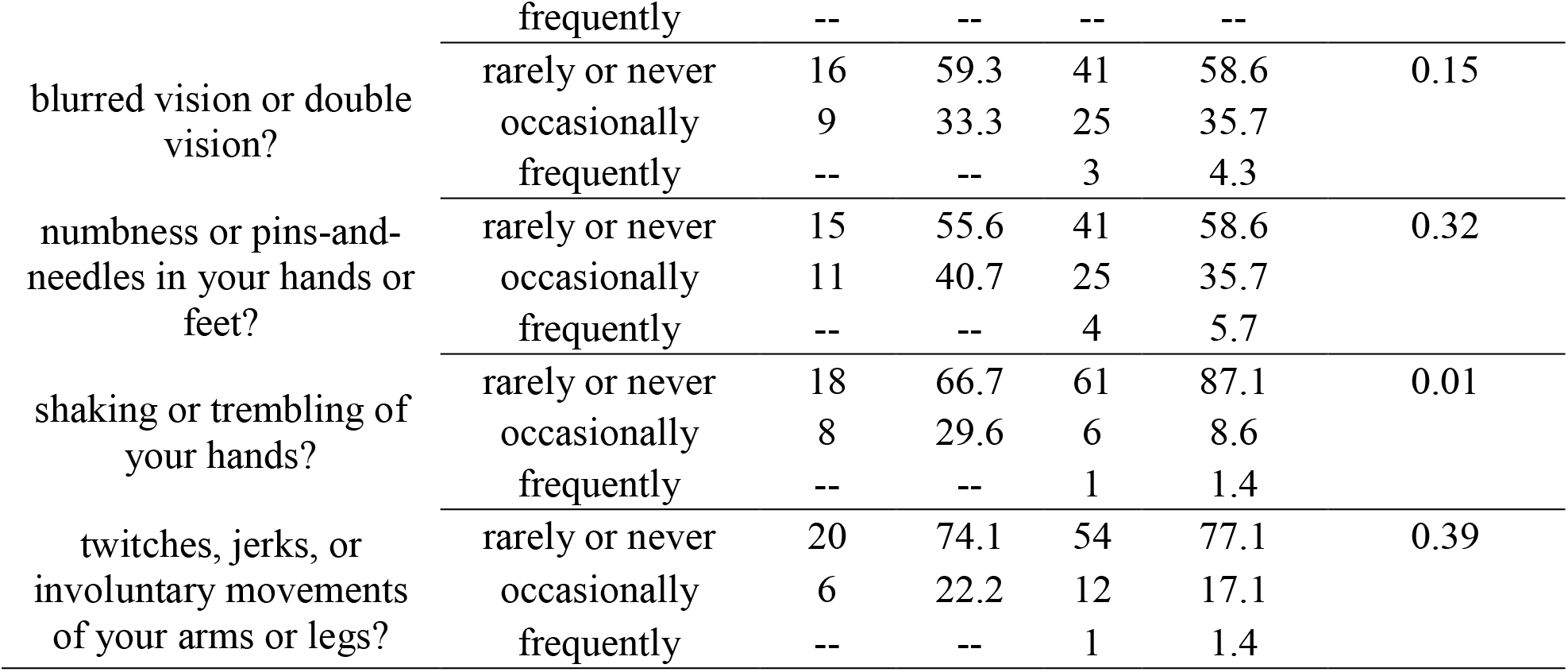
Northern Thailand Participant Self-reported Symptoms by worker category

**Supplemental Table 3.**
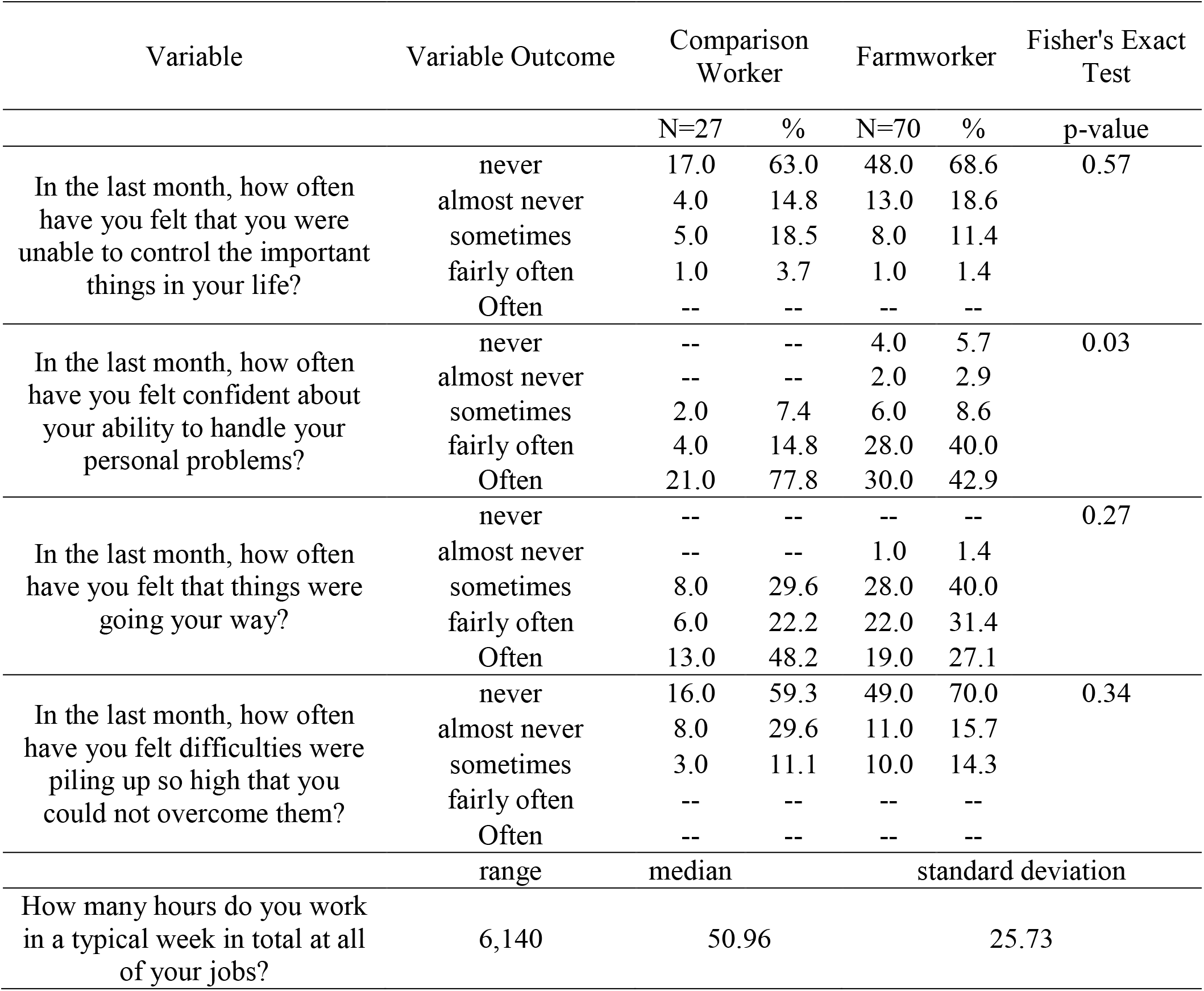
Northern Thailand Participant Stressor Responses 2017, by worker category

| Variable | Variable Outcome | Comparison Worker |  | Farmworker |  | Fisher's Exact Test |
| --- | --- | --- | --- | --- | --- | --- |
|  |  | N=27 | % | N=70 | % | p-value |
| In the last month, how often have you felt that you were unable to control the important things in your life? | never | 17.0 | 63.0 | 48.0 | 68.6 | 0.57 |
|  | almost never | 4.0 | 14.8 | 13.0 | 18.6 |  |
|  | sometimes | 5.0 | 18.5 | 8.0 | 11.4 |  |
|  | fairly often | 1.0 | 3.7 | 1.0 | 1.4 |  |
|  | Often | -- | -- | -- | -- |  |
| In the last month, how often have you felt confident about your ability to handle your personal problems? | never | -- | -- | 4.0 | 5.7 | 0.03 |
|  | almost never | -- | -- | 2.0 | 2.9 |  |
|  | sometimes | 2.0 | 7.4 | 6.0 | 8.6 |  |
|  | fairly often | 4.0 | 14.8 | 28.0 | 40.0 |  |
|  | Often | 21.0 | 77.8 | 30.0 | 42.9 |  |
| In the last month, how often have you felt that things were going your way? | never | -- | -- | -- | -- | 0.27 |
|  | almost never | -- | -- | 1.0 | 1.4 |  |
|  | sometimes | 8.0 | 29.6 | 28.0 | 40.0 |  |
|  | fairly often | 6.0 | 22.2 | 22.0 | 31.4 |  |
|  | Often | 13.0 | 48.2 | 19.0 | 27.1 |  |
| In the last month, how often have you felt difficulties were piling up so high that you could not overcome them? | never | 16.0 | 59.3 | 49.0 | 70.0 | 0.34 |
|  | almost never | 8.0 | 29.6 | 11.0 | 15.7 |  |
|  | sometimes | 3.0 | 11.1 | 10.0 | 14.3 |  |
|  | fairly often | -- | -- | -- | -- |  |
|  | Often | -- | -- | -- | -- |  |
|  | range | median |  | standard deviation |  |  |
| How many hours do you work in a typical week in total at all of your jobs? | 6,140 | 50.96 |  | 25.73 |  |  |

## References

1. Pesticides C. Cholinesterase Testing Protocols for Healthcare Providers. 2010;(2004):22284.

2. Panuwet P, Siriwong W, Prapamontol T, Ryan P barry, Fiedler N, Robson MG, et al. Agricultural Pesticide Management in Thailand: Situation and Population Health Risk. Env sci policy. 2012;17:72–81.

3. Tawatsin A., Thavara, U & Siriyasatien P. Pesticides used in Thailand and toxic effects to human health. Med Res Arch [Internet]. 2015;(3):1–10. Available from: http://www.journals.ke-i.org/index.php/mra/article/view/176

4. Kearns JP, Knappe DRU, Summers RS. Synthetic organic water contaminants in developing communities: an overlooked challenge addressed by adsorption with locally generated char. J Water Sanit Hyg Dev. 2014;4(3):422–36.

5. Thapinta A, Hudak PF. Pesticide use and residual occurrence in Thailand. Environ Monit Assess. 2000;60(1):103–14.

6. Kunstadter P, Prapamontal T, Sirirojn BO, Sontirat A, Tansuhaj A, Khamboonruang C. Pesticide exposures among Hmong farmers in Thailand. Int J Occup Environ Health. 2001;7(4):313–25.

7. Plianbangchang P, Jetiyanon K, Wittaya-areekul S. Pesticide use patterns among smallscale farmers: A case study from phitsanulok, Thailand. Southeast Asian J Trop Med Public Health. 2009;

8. Vasquez C. Illegal Pesticide Trade in Mekong Countries: Pesticide Action Network Asia and the Pacific (PANAP) Sustainable Agriculture and Environment Development Association (SAEDA). 2011;

9. Kaewboonchoo O, Kongtip P, Woskie S. Occupational health and safety for agricultural workers in Thailand: gaps and recommendations, with a focus on pesticide use. New Solut [Internet]. 2015;25(1):102–20. Available from: http://www.scopus.com/inward/record.url?eid=2-s2.0-84955267390&partnerID=tZOtx3y1

10. Panuwet P, Prapamontol T, Chantara S, Thavornyuthikarn P, Montesano MA, Whitehead RD, et al. Concentrations of urinary pesticide metabolites in small-scale farmers in Chiang Mai Province, Thailand. Sci Total Environ [Internet]. 2008;407(1):655–68. Available from: http://dx.doi.org/10.1016/j.scitotenv.2008.08.044

11. Lee WJ, Sandler DP, Blair A, Samanic C, Cross AJ, Alavanja MCR. Pesticide use and colorectal cancer risk in the Agricultural Health Study. Int J cancer [Internet]. 2007;121(2):339–46. Available from: http://www.scopus.com/inward/record.url?eid=2-s2.0-34250329560&partnerID=tZOtx3y1

12. Oddone E, Modonesi C, Gatta G. Occupational exposures and colorectal cancers: A quantitative overview of epidemiological evidence. World J Gastroenterol. 2014;20(35):12431–44.

13. Lee WJ, Hoppin JA, Blair A, Lubin JH, Dosemeci M, Sandler DP, et al. Cancer Incidence among Pesticide Applicators Exposed to Alachlor in the Agricultural Health Study. Am J Epidemiol. 2004;159(4):373–80.

14. Mostafalou S, Abdollahi M. Pesticides and human chronic diseases: Evidences, mechanisms, and perspectives. Vol. 268, Toxicology and Applied Pharmacology. 2013. p. 157–77.

15. Juntarawijit C, Juntarawijit Y. Association between diabetes and pesticides: A casecontrol study among Thai farmers. Environ Health Prev Med. 2018;23(1):1–10.

16. Weiss F, Leuzinger M, Zurbrügg C, Eggen R. Chemical Pollution in Low- and MiddleIncome Countries. Swiss Fed Inst Aquat Sci Technol. 2016;164.

17. Burns KN, Sun K, Fobil JN, Neitzel RL. Heart rate, stress, and occupational noise exposure among electronic waste recycling workers. Int J Environ Res Public Health. 2016;13(1).

18. Ellman GL, Courtney KD, Andres V, Featherstone RM. A new and rapid colorimetric determination of acetylcholinesterase activity. Biochem Pharmacol. 1961;7:88–95.

19. Strelitz J, Engel LS, Keifer MC, Clinic M. Blood acetylcholinesterase and butyrylcholinesterase as biomarkers of cholinesterase depression among pesticide handlers. Occup Env Med. 2014;71(12):842–7.

20. National Institute of Occupational Safety and Health (NIOSH). Organophosphorus Pesticides Method 5600. NIOSH Man Anal Methods. 1993;4(1):419–73.

21. Torres CM, Picó Y, Mañes J. Determination of pesticide residues in fruit and vegetables. J Chromatogr A. 1996;754(1–2):301–31.

22. Gupta RC, Crissman JW. Agricultural Chemicals [Internet]. Third Edit. Haschek and Rousseaux’s Handbook of Toxicologic Pathology. Elsevier; 2013. 1349–1372 p. Available from: http://dx.doi.org/10.1016/B978-0-12-415759-0.00042-X

23. Wilson BW, Arrieta DE, Henderson JD. Monitoring cholinesterases to detect pesticide exposure. Chem Biol Interact. 2005;157–158(November):253–6.

24. Stefanidou M, Athanaselis S, Spiliopoulou H. Butyrylcholinesterase: Biomarker for exposure to organophosphorus insecticides. Intern Med J. 2009;39(1):57–60.

25. Hofmann JN, Keifer MC, De Roos AJ, Fenske RA, Furlong CE, Van Belle G, et al. Occupational determinants of serum cholinesterase inhibition among organophosphate-exposed agricultural pesticide handlers in Washington State. Occup Environ Med. 2010;67(6):375–86.

26. Thetkathuek A, Daniell W. Migrant workers in agriculture: a view from Thailand. J Agromedicine [Internet]. 2016;21(1):106–12. Available from: http://www.tandfonline.com/loi/wagr20

27. Sapbamrer R, Nata S. Health symptoms related to pesticide exposure and agricultural tasks among rice farmers from northern Thailand. Environ Health Prev Med. 2014;19(1):12–20.

28. Andreescu S, Marty JL. Twenty years research in cholinesterase biosensors: From basic research to practical applications. Biomol Eng. 2006;23(1):1–15.

29. Amine A, Mohammadi H, Bourais I, Palleschi G. Enzyme inhibition-based biosensors for food safety and environmental monitoring. Biosens Bioelectron. 2006;21(8):1405–23.

30. García-García CR, Parrón T, Requena M, Alarcón R, Tsatsakis AM, Hernández AF. Occupational pesticide exposure and adverse health effects at the clinical, hematological and biochemical level. Life Sci [Internet]. 2016;145:274–83. Available from: http://dx.doi.org/10.1016/j.lfs.2015.10.013

31. Hu R, Huang X, Huang J, Li Y, Zhang C, Yin Y, et al. Long- and short-term health effects of pesticide exposure: A cohort study from China. PLoS One. 2015;10(6):1–13.

32. Dundar ZD, Ergin M, Koylu R, Ozer R, Cander B, Gunaydin YK. Neutrophil-lymphocyte ratio in patients with pesticide poisoning. J Emerg Med [Internet]. 2014;47(3):286–93. Available from: http://dx.doi.org/10.1016/j.jemermed.2014.01.034

